# Impact of music on running fatigue: Distraction effect from lyrics could further delay running fatigue compared to synchronous effect from tempo

**DOI:** 10.64898/2026.03.09.710701

**Authors:** Megan Dreher, Alexis Terterov, Olivia Feistner, Laura Freiermuth, Polly Schaps, Haley Yeager, Janet Zhang-Lea

**Affiliations:** Department of Human Physiology, Gonzaga University, Spokane, Washington, United States of America; Department of Sport and Exercise Sciences, Durham University, Durham, England; Department of Anthropology, Durham University, Durham, England; Department of Human Physiology, University of Oregon, Eugene, Oregon, United States of America

**Author notes:** Corresponding Author: Janet Zhang-Lea, Address: 1484 University Street, Gerlinger Annex 171, Eugene, Oregon, USA.

**Keywords:** Running biomechanics, Running fatigue, Music lyrics, Tempo

## Abstract

Motivational music has been shown to improve running performance through delaying fatigue and increasing run duration. Previous studies have highlighted the effect of music tempo, that matching tempo to the runner’s cadence delays running fatigue. It remains unclear whether the motivational content in music lyrics is also responsible for delaying running fatigue. We designed a cross-sectional study and investigated the effect of tempo and motivational content on running biomechanics, and had participants run at a moderate intensity for up to ten minutes, or until exhaustion. Fifteen adults (age=20.9±1.3 years, weight=71.2±12.1 kg, height=174.7±11.0 cm) participated. Participants finished three trials, starting with running without any stimulus as a baseline trial, and ran with a visual metronome that flashed at a rate that matched their running cadence in the visual stimulus trial (VST). In the visual-auditory stimulus trial (VAST), participants ran with the visual metronome (as described in VST) while listening to a non-rhythmic motivational speech. We recorded run duration, perceived exertion, center of pressure sway during standing before and after each trial, and measured trunk acceleration to obtain root-mean-square (RMS) of acceleration during each minute of the run. Compared to baseline, participants reduced perceived exertion by 0.87 and 0.85 rating during the VST and VAST, respectively, though these changes did not reach significance (p=0.05). Stimulus affected the RMS of acceleration in anterior-posterior (p=0.011), vertical (p=0.008), and resultant directions (p=0.006). Our linear mixed effect model suggested that compared VST, VAST further lowered RMS of acceleration by 0.026g (anterior-posterior), 0.028g (vertical), and 0.036g (resultant). Our results showed that motivational content played an important role in lowering RMS of trunk acceleration, with the potential to delay running-induced fatigue. To maximize the effect of music on running performance, runners should listen to music that they find motivational and that is close to their natural running cadence.

## Introduction

Motivational music has been found to elicit positive emotional responses in runners, to improve run performance, and to extend run duration [1,2]. The effect of music in impeding running fatigue has been attributed to the tempo and the motivational lyrics of music [3]. Music tempo could generate an *synchronous effect*, which occurs when runners temporally regulate their running cadence to the tempo of music [3]. Synchronicity induced by the tempo of music is shown to lower oxygen consumption in runners, indicating a reduced muscle activation and improved preservation of metabolic energy [4,5]. Synchronous music has been found to reduce perceived exertion and to increase run duration by up to 15% [6–8]. Prior studies show that runners adjust their running cadence, which is usually between 130 and 200 steps per minute [9], in accordance with a music tempo within 2.5-10% of their natural cadence to increase their running speed [9–11]. However, this adaptation may be influenced by the instructions of researchers rather than a natural entrainment of running cadence to music tempo [11].

On the other hand, motivational content stemming from music lyrics may divert attention from feelings of fatigue while running [3,12], and therefore could extend run duration through *distraction effect*, as described in the attentional processing mechanism [3,12]. The distraction effect of music is harder to quantify since all components of music can be argued as a distraction to the runners from feeling fatigued, which includes motivational lyrics, tempo, melody, etc. Previous studies aimed to assess the distraction effect of music, but did not separately study the effect of music tempo and the effect of motivational lyrics on running fatigue and running biomechanics [Deforche et al., 2015]. Therefore, it is unclear whether the previously reported effect of music in delaying fatigue is solely due to the synchronous effect of music tempo or it’s a combination of tempo and motivational lyrics of music.

In this study, we aim to understand the effect of music tempo and the combined effect of tempo and motivational lyrics on running fatigue and running biomechanics. In other words, we aim to learn if motivational lyrics will bring in additional benefits in delaying running fatigue and affecting fatigue-related running biomechanics variables. We will use measures of run duration, perceived exertion, and fatigue-related biomechanics variables to assess the onset of running-induced fatigue [13–15]. Perceived exertion is a subjective, internal measure of peripheral discomfort, that is used to assess running fatigue [13]. Additionally, after running, runners typically present with increased fatigue, increased ventilation or disturbances in vestibular, visual, and proprioceptive inputs [14], which affects the postural control during standing and center of mass osculation during running. Therefore, several biomechanical variables, such as standing center of pressure (CoP) sway [14] and deviation of trunk acceleration [15] during running have been used to assess running-induced fatigue.

To our knowledge, prior research investigating the effects of music on running fatigue has not distinguished between the rhythmic and distraction effects of music, making it difficult to discern which mechanism has the greatest impact on delaying running fatigue. Our study sought to comparatively investigate the rhythmic and distraction effects of music on run duration, perceived exertion, and running biomechanics in runners. We dissected music into tempo and motivational lyrics. We utilized a flashing visual metronome to simulate the rhythmic effect of music and control running cadence, and we hypothesized that compared to running without a stimulus, runners would prolong their run, experience less perceived exertion, reduce deviations in their root-mean-squares (RMS) of trunk acceleration, and reduce their standing CoP sway when matching their running cadence with the flashing of the visual metronome. To simulate the distraction effect of music, we utilized a non-rhythmic motivational speech that was unfamiliar to participants, and we hypothesized that compared to running with only the visual metronome, runners would further prolong their run, experience less perceived exertion, reduce RMS of trunk acceleration, and further reduce standing CoP sway after running when listening to the motivational speech.

## Methods

### Participants

We conducted a prior sample size estimation using G*Power [16] based on previously published effect size values. Since to our best knowledge, no study has reported the effect of music on root-means-squares (RMS) of trunk acceleration, or center of pressure, we used previous publications reporting the effect of music on oxygen consumption (Cohen’s f = 1.03) [5], electromyographic fatigue threshold (Cohen’s f = 1.94) and lower limb muscle maximal power output (Cohen’s f = 1.66) [17]. We set the power at 0.8, and the type I error rate at 5%, and our estimate showed a sample size between 4-8 participants will be sufficient to reach the targeted statistical power.

Fifteen young adults (7 males, 8 females, age= 20.9 ± 1.3 years, body weight= 71.2 ± 12.1 kg, body height= 174.7 ± 11.0 cm) participated in this study. Participants were over 18 years old and self-reported previously using a treadmill for moderate-intensity exercise. All participants were free of cardiovascular, neurological, and musculoskeletal conditions. Additional exclusion criteria included untreated vision or hearing loss as well as a history of epilepsy or light sensitivity. This study was approved by Institutional Review Board at Gonzaga University, and the recruitment period started June 16, 2023, and ended on May 15, 2024. Written informed consents were obtained from all participants prior to their participation.

### Experimental Protocol

All participants first completed an overground sprinting trial for us to obtain their maximum speed. Participants accelerated for 20 meters before running through two speed gates (DASHR, Lincoln, Nebraska) spaced 3 meters apart. We recorded the average velocity within the 3-meter distance between the two speed gates as the participant’s maximal, reported in m/s. Participants were given three times to sprint through the speed gates, we took the fastest speed among the three trials as the individual’s maximum speed. We used 55% of each participant’s maximum speed as their testing speed.

Participants then completed baseline, visual stimulus (VST), and visual-auditory stimulus (VAST) treadmill (Woodway, Waukesha, Wisconsin) running trials at testing speed on separate days with at least five days between trials (Figure 1). All participants completed a structure, dynamic warm-up led by one of our experimenters prior to each run. When beginning each run, we hid the speed and running time from participants and increased the speed by 0.45 m/s every 3 seconds until the testing speed was reached. A Polar Heart Rate Chest Strap (Polar, Kempele, Finland) was fixed to a participant’s sternum monitoring heart rate (HR) continuously and maximum HR was defined as the sum of 220 minus a participant’s age [18]. Participants were asked to verbally rate themselves on the Borg Rating of Perceived Exertion Scale (RPE) as a measure of perceived fatigue [13]. HR and RPE were recorded every minute. Trials were terminated when (A) maximum HR was reached, (B) a score of 17 was reported on the RPE, (C) the treadmill safety clip was pulled, or (D) the trial lasted for ten minutes. Participants were blinded to the duration of each trial. Participants wore a triaxial accelerometer (±16g, Delsys, Natick, MA) on the back of their L1 vertebrae, secured by athletic tape, throughout the duration of each trial. The accelerometer measured trunk acceleration along the medio-lateral, vertical, and anterior-posterior axes at 148 Hz. Immediately before and after each running trial, Participants stood hip-width apart on a Kistler force plate (9287BA, Kistler Group, Novi, Michigan) for 10 seconds and we recorded CoP trajectory at 1000 Hz using BioWare (Kistler Group, Alberta, Canada).

**Figure 1.**
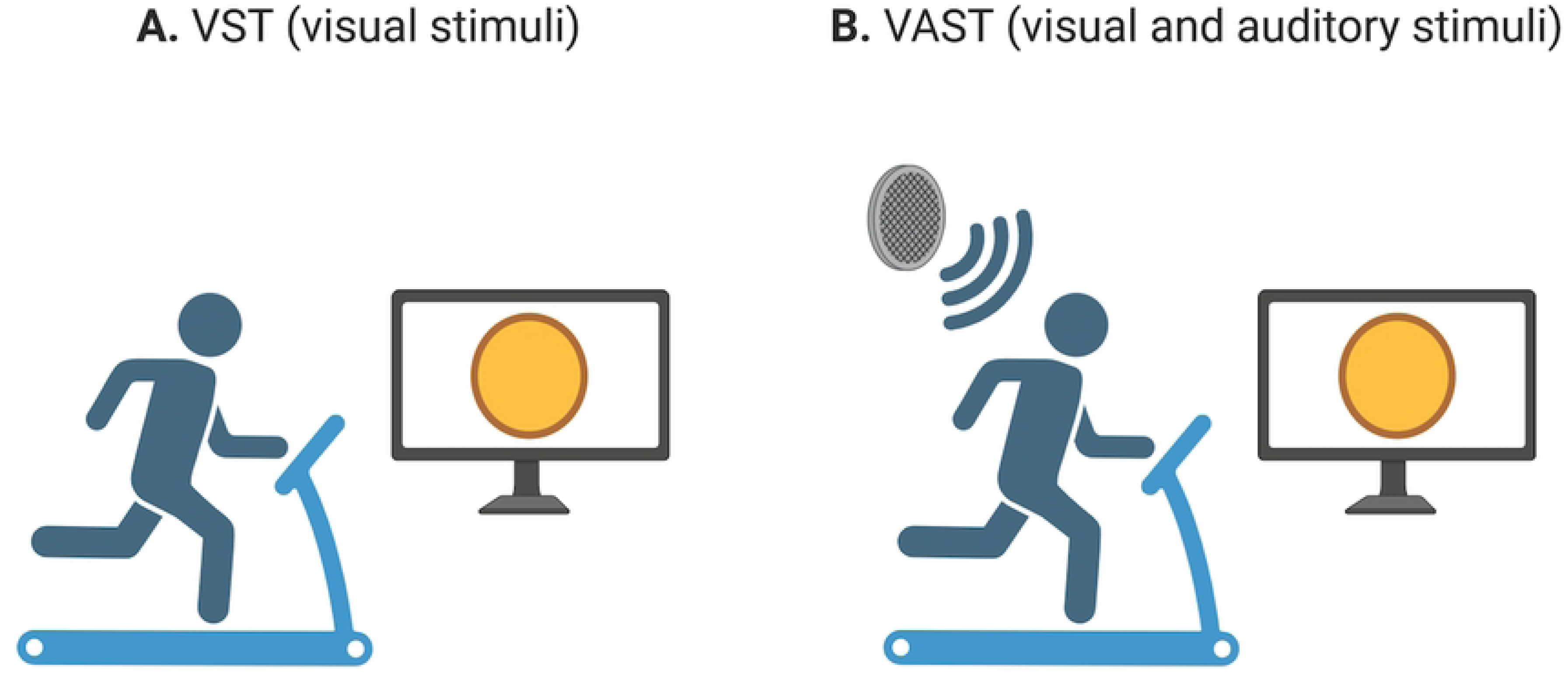
Overview of experimental setup for visual stimulus (VST) and visual-auditory stimulus trials (VAST). Participants completed all three treadmill running trials at 55% of their maximum speed, and ran with no feedback during baseline trial. In VST (A) and VAST (B), participants ran with a visual metronome shown as a flashing yellow circle on a screen positioned in front of the treadmill at eye level. Additional auditory stimulus was provided during VAST via a speaker positioned on the treadmill at the same spot for every trial (B).

All participants completed baseline trial first and completed VST and VAST in a randomized order. During baseline running, we used a smartphone (iPhone 12, Apple, Cupertino, CA) to record the first ten seconds of the participant running at test speed at a 60 Hz frame rate. This footage was used in conjunction with a mobile phone application (Metronome, Soundbrenner, Hong Kong, China) to calculate cadence as the total number of steps a participant took per minute. We calibrated a custom-written visual metronome (MATLAB, R2022B, MathWorks, Natick, Massachusetts), to flash a yellow circle at the same frequency as the participant’s cadence obtained at baseline. During VST, the visual metronome was displayed on an eye-level screen in front of the treadmill and all participants were instructed to couple their running steps with the flashes (Figure. 1A). During VAST, participants were instructed to couple their running steps with the same visual metronome while listening to a 10-minute audio recording (Figure 1B), which did not include any melody or rhythmic stimulation [19]. Using a visual metronome in this study allowed the participants to use this visual cue to guide their cadence while being able to receive motivational content through the audio recording that is non-rhythmic. We played the audio recording via a speaker placed on the treadmill, and the recording was played at a predetermined volume and kept the same for all trials and all participants.

The motivational speech we chose for this study was delivered by former American basketball coach, Jim Valvano, on March 4, 1993, shortly before he passed away due to cancer. We chose this speech for many reasons. First, the length of the speech is 10 minutes 16 seconds, which aligns with the maximal run duration of our study (10 minutes). This is important because it allows our participants to be able to follow the paragraphs of content delivered by the speaker to the maximal level. Second, the theme of the speech aligns closely with sports and exercise. The speaker, Jim Valvano, was a basketball coach, and the speech was delivered when receiving the Excellence in Sports Performance Yearly award. In the speech, he openly spoke about having terminal cancer and showcases his resilience in the face of mortality, and the theme “Don’t give up. Don’t ever give up” runs through the entire speech. Third, this speech does not have any music component (e.g. rhythm, drumbeats) in the background, reducing confounding factors that may complicate the findings of our study.

### Data Analysis

We first verified the cadence each participant ran with during each stimulus trial with the flash rate set at the visual metronome. We used a custom written program to identify ten peaks in the vertical acceleration data for ten consecutive steps in the middle of the trial. The time intervals between each step were used to calculate cadence for the specific trial.

We filtered CoP data and trunk acceleration using a 4^th^ order Butterworth low-pass filter with a cutoff frequency at 30 and 50 Hz respectively. We then performed a principal component analysis on CoP data and fitted a 95% confidence interval ellipse to CoP trajectory during standing. We defined the major axis length of the confidence ellipse as the frontal sway, and the minor axis length as the sagittal sway. For the trunk acceleration data, we first adjusted for gravitational acceleration for each axis and segmented the continuous acceleration data into 1-minute intervals. For participants with a final segment shorter than 1-minute, we reserved the data points if there were more than 10 seconds of running in the segment. We used a previously reported method to calculate root-mean-squares (RMS) of the trunk acceleration data [15]. Specifically, for each segment of data, we calculated the resultant acceleration and calculated the RMS of acceleration in the medio-lateral (ML_RMS_), anterior-posterior (AP_RMS_), vertical (VT_RMS_), as well as a resultant (RES_RMS_) directions [15]. We then divided the RMS of acceleration along each axis with the resultant RMS of acceleration to get the ratio of acceleration (ML_ratio_, AP_ratio_, VT_ratio_).

### Statistical Analysis

The Shapiro-Wilk test was used to test for data normality. If the normality assumption was met, we used one-way repeated measures ANOVAs to verify if the participant’s cadence during VST and VAST match with rate of visual metronome flashes. We used one-way repeated measures ANOVAs to compare the time to fatigue across all trials. We constructed several linear mixed models to compare the effects of running time (abbreviated as RT: 1^st^, 2^nd^, 3^rd^, etc. minute of running), stimulus condition (abbreviated as STIM: baseline, VST, VAST), and the interaction effect between RT and STIM on RPE and acceleration-related variables (RMS and RMS ratio). We constructed linear mixed models to compare the effects of trial completion (pre vs. post) and STIM on standing sagittal and frontal sway of CoP. All statistical tests were performed using MATLAB, with level of significance set at 0.05. If post-hoc analysis is needed, we conducted all post-hoc analysis in G*Power [16].

## Results

One participant stopped their VST by accident, and their data were excluded from all further analysis. Two participants were unable to record CoP data post-trial. The accelerometer detached from four participants before they reached fatigue, and their data were excluded from analysis of time to fatigue, cadence, RPE, and all trunk acceleration variables. Among all 15 participants, 12 were included in analysis of CoP, and 10 were included in the analysis of RPE and acceleration-related variables. Participants ran at similar cadences (F (2,27) =0.027, p=0.97) in baseline (171.2± 10.4 SPM), VST (171.6± 12.7 SPM), and VAST (171.3± 13.4 SPM), and with similar run duration (F (2,27) =0.038, p=0.96, baseline: 350.26 ± 192.80 s, VST: 371.01±189.30 s, VAST: 369.60± 185.80 s). Participants that stopped prior to ten-minutes also presented similar run duration across the three trials (F (2,18) =0.15, p=0.86, baseline: 241.21 ± 97.33 s, VST: 269.57 ± 117.17 s, VAST: 265.20 ± 96.90 s).

### Rate of perceived exertion when running with varied stimulus conditions

Run time had a significant effect on RPE (F=91.13, p <0.001), such that RPE increased by 0.585 for every minute increase in running time (p<0.001), and STIM trended towards having an impact on RPE (F=3.761, p=0.05) but did not reach significance (Figure 2). Linear mixed models showed that compared to baseline, participants reported RPE values were 0.87 less during VST (p=0.05) and 0.85 less during VAST (p=0.05). RPEs between VST and VAST were not different (p=0.97). We did not find an interaction effect between RT and STIM on RPE, F=0.93, p=0.34).

**Figure 2.**
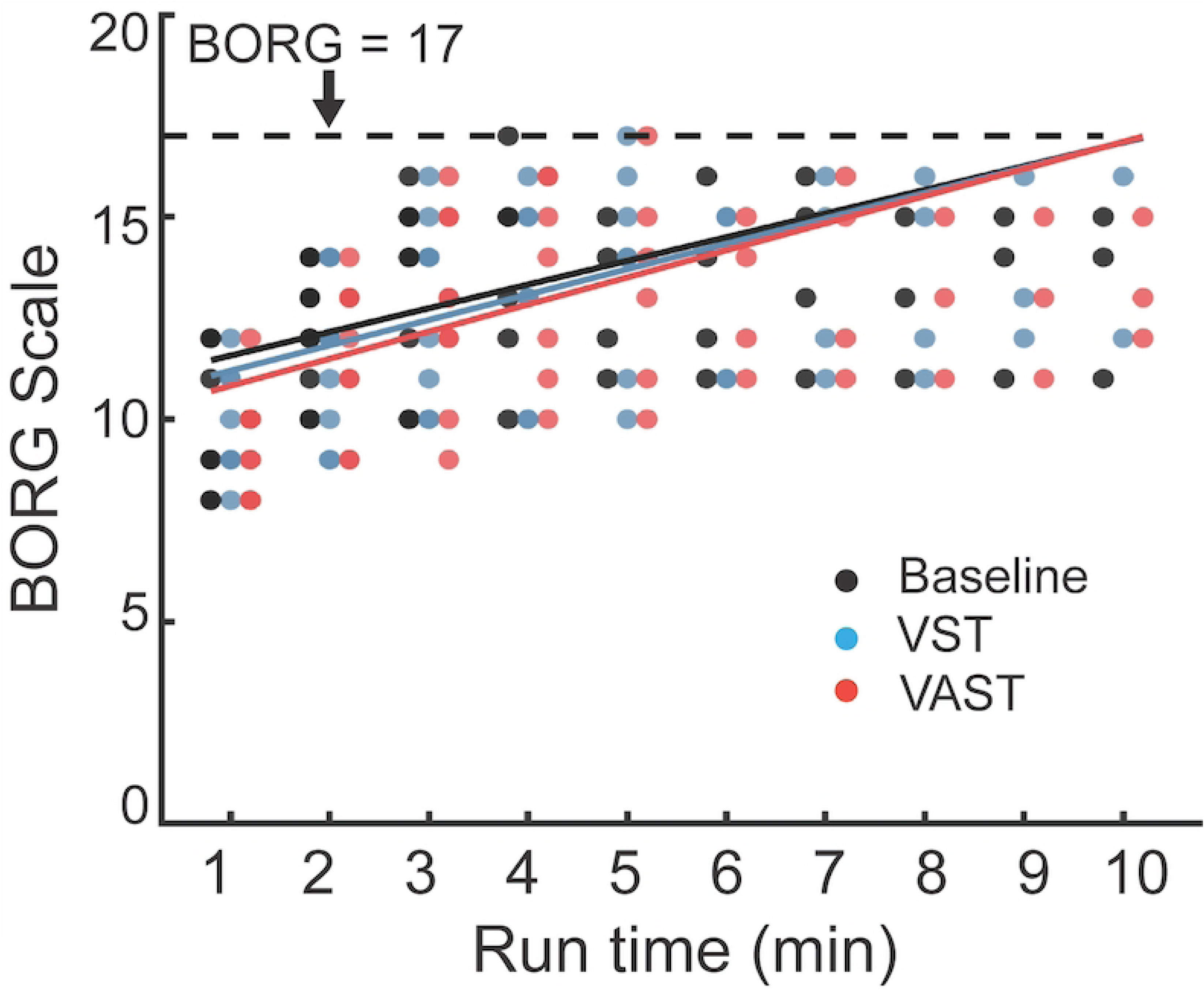
Effect of run time and stimulation on perceived fatigue, measured using the BORG scale. Each individual dot represents data from one participant. The three solid lines represents regression model generated based on linear mixed effect model for baseline (black), VST (blue), and VAST (red) trials. The dotted line represents BORG scale reaches 17, which is one of the criteria to stop the trial.

### Static CoP sway before and after running with varied stimulus conditions

When controlling for STIM and interaction effect, compared to pre-running, CoP_frontal_ increased by 0.026 m^2^ (p=0.013) and CoP_sagittal_ increased by 0.017 m^2^ (p=0.001) after completion of running trials (Table 1). When controlling for trial completion and interaction effect, STIM did not affect static CoP sway along the frontal (p= 0.56) or sagittal axis (p=0.53) (Figure 3). We did not find any interaction effects between trial completion and stimulation for either CoP_frontal_ (p=0.833) or CoP_sagittal_ (p=0.416).

**Table 1.**
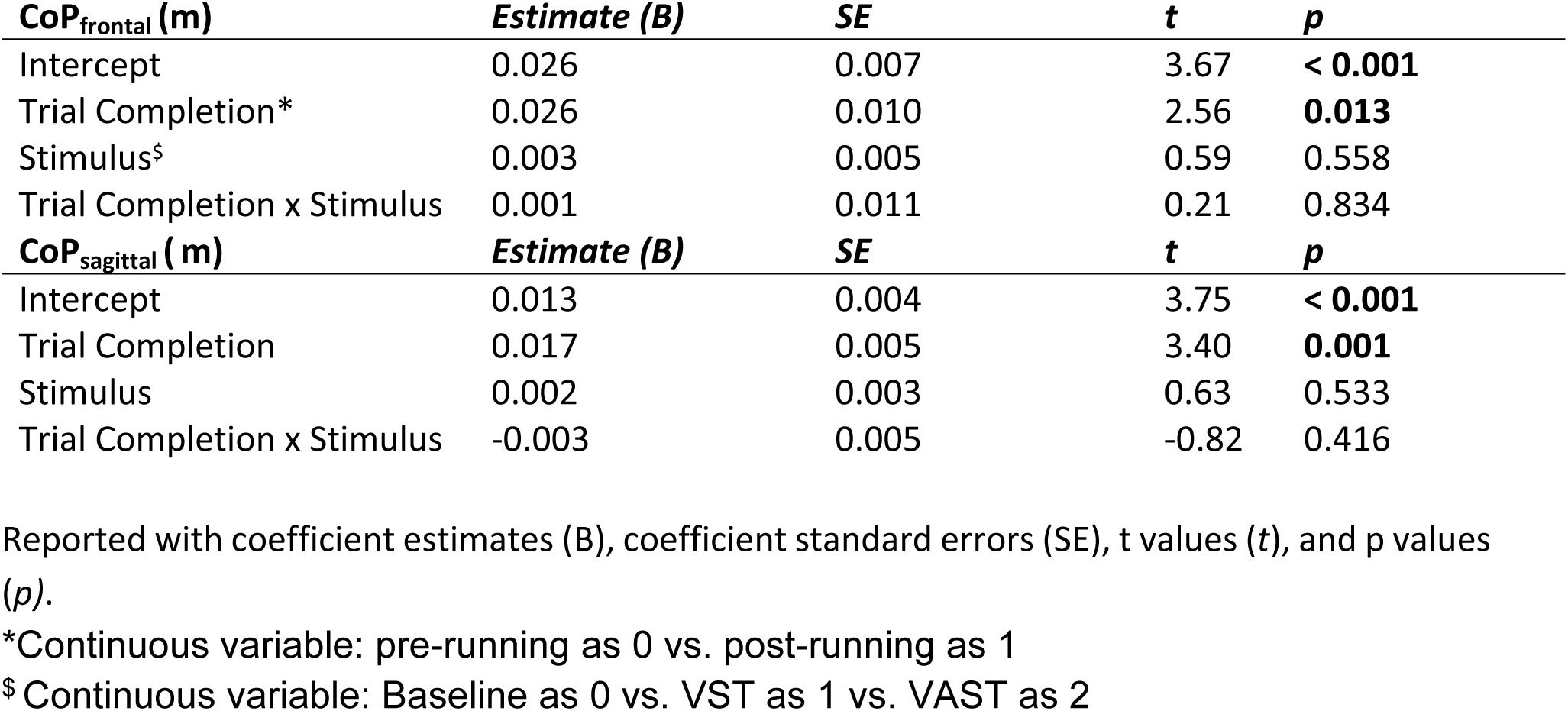
Linear mixed model parameters for fixed effects of Trial Completion (pre-running vs. post-running) and Stimulus (baseline, VST, VAST) on static CoP sway along the frontal and sagittal axes.

**Figure 3.**
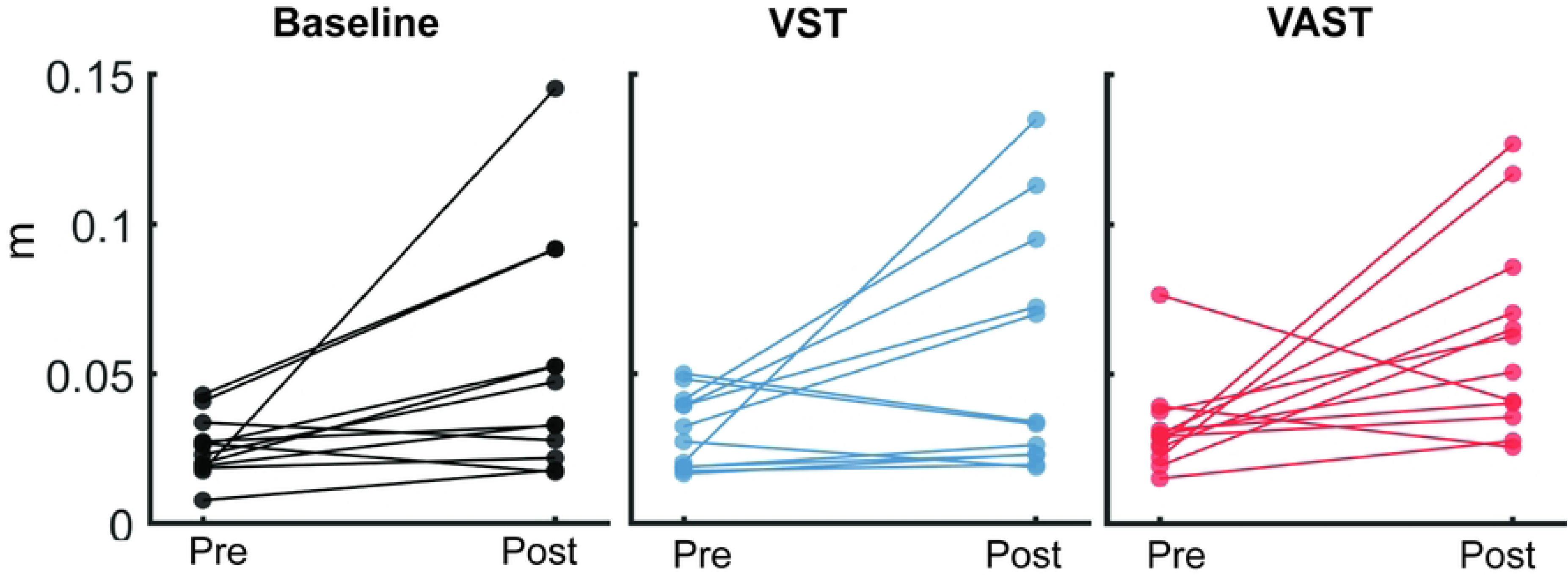
Static center of pressure sway increased after trial completion but did not vary between trials. Boxplots representing pre- and post-trial static CoP sway data for baseline (white), VST (gray), and VAST (black). All data is reported in the unit of meters.

### RMS of acceleration and ratio of acceleration when running with varied stimulus conditions

When controlling for STIM and interaction effect, running time affected the RMS of acceleration in the anterior-posterior (p=0.037), medio-lateral (p<0.001) and resultant directions (p=0.001), but not in the vertical (p=0.35) direction (Figure 4). Specifically, AP_RMS_ increased by 0.006 g, ML_RMS_ increased by 0.016 g, and RES_RMS_ increased by 0.011 g (Table 2) for every minute of running. When controlling for running time and interaction effect, STIM affected the RMS of acceleration in anterior-posterior (p=0.011), vertical (p=0.008), and resultant directions (p=0.006), but not in the medio-lateral direction (p=0.79, Figure 4). Based on the linear mixed effect model, adding one type of stimulation lowers AP_RMS_ by 0.026 g, VT_RMS_ by 0.028 g, and RES_RMS_ by 0.036 g (Table 2). We did not find any interaction effect between STIM and run time.

**Figure 4.**
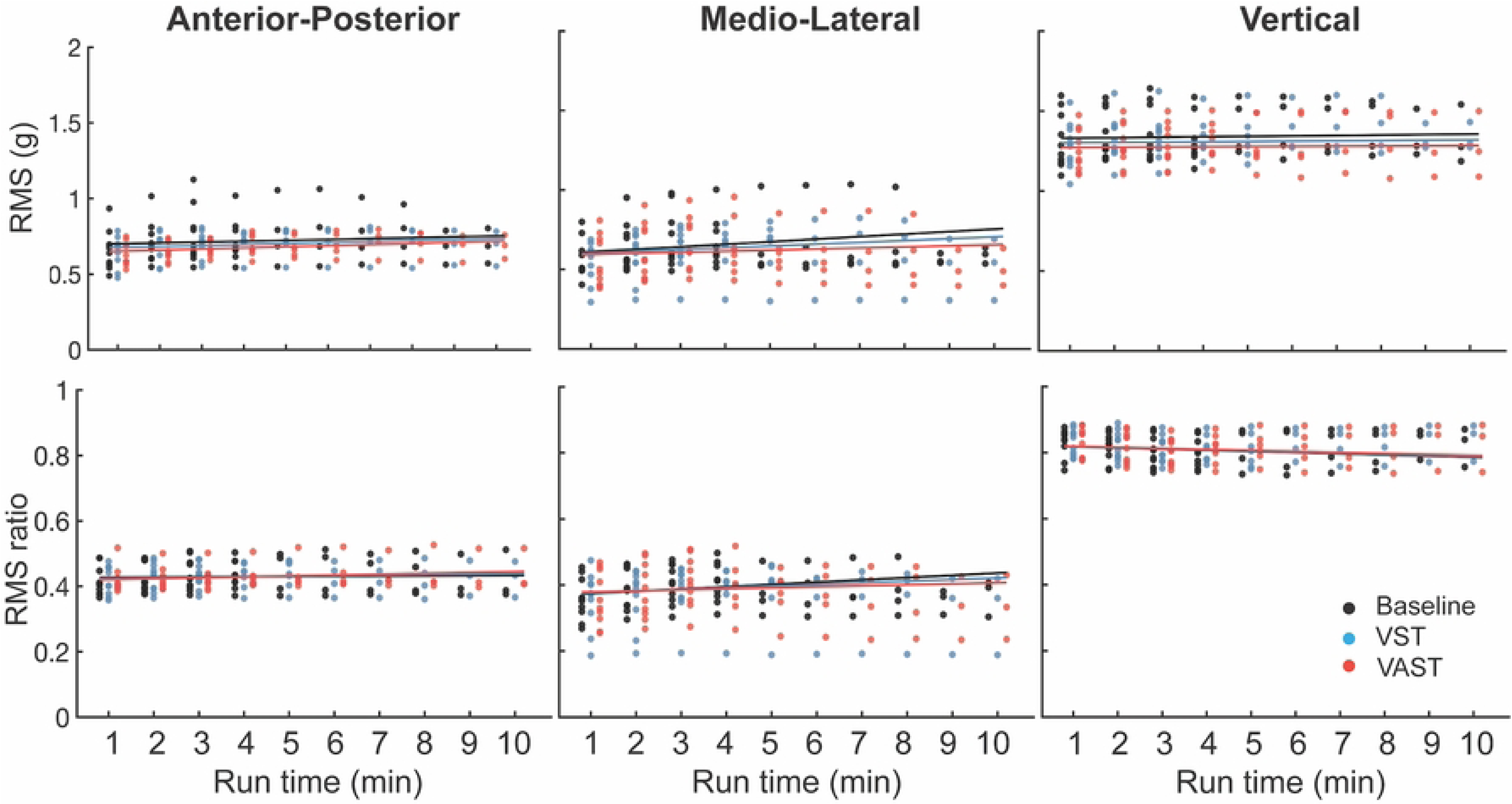
RMS of trunk acceleration along the medio-lateral, anterior-posterior, and vertical directions was lowest for VAST. RMS of trunk acceleration reported via boxplot for baseline (white), VST (grey), and VAST (black) for each minute of running time.

**Table 2.** Linear mixed model parameters for fixed effects of Running Time (min) and Stimulus (baseline, VST, VAST) on RMS of Acceleration.

| **$AP_{RMS}$ (G)** | ***Estimate (B)*** | ***SE*** | ***t*** | ***p*** |
| --- | --- | --- | --- | --- |
| Intercept | 0.694 | 0.030 | 23.10 | **< 0.001** |
| Run Time (Minutes) | 0.006 | 0.003 | 2.10 | **0.037** |
| Stimulus* | -0.026 | 0.010 | -2.58 | **0.011** |
| Run Time x Stimulus | <0.001 | 0.002 | 0.40 | 0.688 |
| **$ML_{RMS}$ (G)** | ***Estimate (B)*** | ***SE*** | ***t*** | ***p*** |
| Intercept | 0.592 | 0.046 | 12.93 | **< 0.001** |
| Run Time (Minutes) | 0.016 | 0.003 | 4.56 | **< 0.001** |
| Stimulus | -0.003 | 0.025 | -0.26 | 0.794 |
| Run Time x Stimulus | -0.004 | 0.024 | -1.79 | 0.076 |
| **$VT_{RMS}$ (G)** | ***Estimate (B)*** | ***SE*** | ***t*** | ***p*** |
| Intercept | 1.327 | 0.041 | 32.57 | **< 0.001** |
| Run Time (Minutes) | 0.003 | 0.003 | 0.94 | 0.347 |
| Stimulus | -0.028 | 0.011 | -2.69 | **0.008** |
| Run Time x Stimulus | <-0.001 | 0.002 | -0.36 | 0.722 |
| **$RES_{RMS}$ (G)** | ***Estimate (B)*** | ***SE*** | ***t*** | ***p*** |
| Intercept | 1.617 | 0.051 | 31.86 | **< 0.001** |
| Run Time (Minutes) | 0.011 | 0.003 | 3.049 | **0.003** |
| Stimulus | -0.036 | 0.013 | -2.78 | **0.006** |
| Run Time x Stimulus | -0.002 | 0.003 | -0.72 | 0.475 |
| **$AP_{ratio}$** | ***Estimate (B)*** | ***SE*** | ***t*** | ***p*** |
| Intercept | 0.426 | 0.011 | 38.60 | **< 0.001** |
| Run Time (Minutes) | <0.001 | 0.001 | 0.58 | 0.560 |
| Stimulus | -0.005 | 0.004 | -1.10 | 0.274 |
| Run Time x Stimulus | 0.008 | <0.001 | 1.22 | 0.225 |
| **ML<sub>ratio</sub>** | **Estimate (B)** | **SE** | **t** | **p** |
| Intercept | 0.366 | 0.023 | 16.23 | **< 0.001** |
| Run Time (Minutes) | 0.007 | 0.002 | 4.00 | **< 0.001** |
| Stimulus | 0.005 | 0.006 | 0.79 | 0.428 |
| Run Time x Stimulus | - 0.002 | 0.001 | -1.55 | 0.123 |
| **VT<sub>ratio</sub>** | **Estimate (B)** | **SE** | **t** | **p** |
| Intercept | 0.822 | 0.013 | 63.21 | **< 0.001** |
| Run Time (Minutes) | -0.003 | 0.001 | -3.32 | **0.001** |
| Stimulus | <0.001 | 0.003 | 0.006 | 0.995 |
| Run Time x Stimulus | <0.001 | <0.001 | 0.32 | 0.749 |

Reported with coefficient estimates (B), coefficient standard errors (SE), t values (t), and p values (p).
\* Continuous variable: baseline as 0 vs. VST as 1 vs. VAST as 2

When controlling for STIM and interaction effect, running time affected the ratio of RMS of acceleration in medio-lateral (p < 0.001) and vertical (p= 0.001) directions, but not anterior-posterior (p= 0756). Specifically, ML_ratio_ increased by 0.007 (p <0.001) and VT_ratio_ decreased by 0.003 (p= 0.001) (Table 2) for every minute running. When controlling for running time and interaction effect, STIM did not affect the ratio of RMS of acceleration in the anterior-posterior (p=0.304), medio-lateral (p= 0.266), or vertical (p= 0.789) directions. Running time did not interact with STIM on any of the directions (p=0.225 for AP_ratio_, p=0.123 for ML_ratio_, p=0.749 for VT_ratio_).

## Discussion

Through measures of run duration, perceived exertion, and fatigue-related biomechanics variables, this study investigated the effects of both synchronous beat and motivational content on running fatigue. Participants ran on the treadmill at 55% of their maximal sprinting speed for ten minutes, or until exhaustion, with no stimulus, with a visual metronome to simulate synchronous beat, and with a non-rhythmic audio recording to simulate motivational content. The addition of the visual metronome to emulate the rhythmic effect was responsible for improvements in perceived exertion compared to baseline, while the combined effect of motivational content and the visual metronome was responsible for reductions in fatigue-related biomechanics compared to baseline.

### Rhythmic effect from the visual metronome is responsible for reduced perceived exertion compared to baseline

We partially accepted our first hypothesis, that participants’ RPE trended towards being greater at baseline compared to both VST and VAST, but we did not find any difference in RPE between VST and VAST. While our study used visual metronome to provide the rhythmic effect of beats, previous research studies using auditory beat stimulation have reported similar results [4,8,20]. One major reason could be that in both our study and prior studies, the frequency of the beat were matched with the runner’s cadence, amplifying the effect of synchronous response [4,8,20]. Some prior studies have proposed that using a metronome in running enabled runners to anticipate their next step and to maintain their optimal cadence when it would otherwise slow down due to fatigue [8,12]. Additionally, prior studies also report that runners can spontaneously entrain their cadence to music with a tempo greater than 120 bpm [21], and some have suggested that the rhythmic effect lowers RPE when the tempo of the music surpasses 120-bpm [22,23]. Since the metronome frequencies were high in our study (171.2±10.4 bpm), it could contribute to the effect of rhythmic effect.

We found similar RPE between VST and VAST (p=0.97), and we therefore suggest that the rhythmic effect of music simulated via the visual metronome is more responsible for a lower RPE compared to distraction effect of music provided via motivational content. Such results are in conflict with some previous studies suggesting the distraction effect of music created by motivational content is also responsible for reductions in RPE [1,24]. One possible explanation is that the previous studies allowed participants to self-select their preferred motivational music, while all our participants listened to the same motivational speech. Listening to the same speech controlled the content that was delivered to the runners, but it is possible that not all participants were similarly engaged with the motivational speech. Our study found that run duration and CoP sway during standing did not differ across trials, indicating that neither the rhythmic effect nor distraction effect improve this aspect of running performance. In our study, only seven participants reached our definition of fatigue when each trial stopped, which limits our interpretation of the effect of stimulation on run duration.

The rhythmic effect of music found in our study aligns with the synchronous response mechanism proposed by Karageorghis and Priest [3]. The synchronous response, more commonly acknowledged as a “rhythm response”, occurs when runners temporally regulate their movement patterns to the tempo, or speed, of music [3]. The synchronicity between music tempo and running cadence has ergogenic implications, as it lowers oxygen consumption in runners [5]. This ergogenic effect is attributed to an increase in neuromuscular metabolic efficiency, meaning that less muscle force is needed to maintain movement patterns when synchronously running with a familiar and regular tempo [4,12,21].

### Motivational content and rhythmic effect together reduced RMS of trunk acceleration compared to baseline

We partially accepted our second hypothesis. Our results suggested that VAST did not further increase run duration or lower RPE values compared to VST. However, we found that the type of stimulation had an impact on RMS of trunk acceleration in the anterior-posterior, vertical, and resultant directions. Specifically, when controlling for time and interaction effect, running with VAST lowers RMS of trunk in the previously mentioned directions more compared to running under VST baseline conditions. An increase in the RMS of acceleration has previously been attributed to running fatigue, as it is suggested to result from the decrease in the knee flexion angle of a runner [25] during fatigued status. Thus, the lower RMS of acceleration observed in each of these directions in VAST compared to baseline suggests that the combined use of a visual metronome and motivational content reduced running fatigue.

Our further pairwise comparison suggested that RMS of trunk acceleration trended towards reduction when comparing VST to baseline, but such difference did not reach significance (p>0.064 for all directions). This result suggests that combining motivational content with the visual metronome best reduced running fatigue, as opposed to findings that credit only the rhythmic effect based on lower measures of RPE, less time spent running, and greater distance traveled, but not the distraction effect from motivational content [7,8]. Such discrepancy in our findings could be due to the fact that runners were exposed to different music pieces in order to match the music beats to their cadence in previous studies. In our study, we controlled the motivational content that runners listened to by separating rhythmic effect and distraction effect. Our findings suggested that distraction effect on its own has impact in delaying running-related fatigue. One proposed explanation is that when exposed to external distraction, runners shift from focusing on internal behavior, such as stride length and step frequency, to focusing on external environment, such as breathing or external environment [26,27]. There have been studies reporting benefits of external focus on attention in endurance sports, including a better running economy, an increased VO_2max_, which are all potentially contribute to a delayed fatigue in endurance running [26,28,29].

Limitations of this study included that all runners listened to one identical motivational content. Although this eliminates the variance of motivational content as a confounding factor, this brings the issue that all runners may not be similarly engaged to the content. Another limitation is that our study quantified fatigue using one accelerometer attached on the trunk, which is considered to associate with running kinematic changes that occur during running fatigue. However, we did not directly measure running kinematics variables in this study. To ensure the safety of the participants and to facilitate participant recruitment and retention, we limited the maximum running duration to 10 minutes, and we expected our participants to reach fatigue prior to 10 minutes. While majority of our participants stopped prior to the 10-minute time mark, there were three participants able to finish all 10 minutes of running, which limits our understanding on the effect of stimulations on run duration.

## Conclusions

Our study found that the distraction effect of music showed additional benefit in delaying running-related fatigue compared to rhythmic effect only condition. This is based on our finding that runners presented lower RMS of trunk acceleration when presented with both rhythmic and distraction effects, compared to presented with rhythmic effect alone. Based on our findings, if runners have access to music, we suggest runners listen to music with motivational lyrics but that also has a tempo within 2.5% of their natural running cadence. In situations when runners are not allowed to listen to music, or when music is not available, motivational talks could also delay running-related fatigue even if they are non-rhythmic and do not align with the runner’s cadence.

## Acknowledgements

The project proposal received undergraduate research award ($992.00) from American College of Sports Medicine Northwest Chapter in 2024 to support part of the project expenses. The funders had no role in study design, data collection and analysis, decision to publish, or preparation of the manuscript.

## Supporting information

**S1. Acceleration and RPE data**

**S2. CoP data**

**S3. Script:** MATLAB script for statistical analysis

## References

1. Hutchinson JC, Jones L, Vitti SN, Moore A, Dalton PC, O’Neil BJ. The influence of self-selected music on affect-regulated exercise intensity and remembered pleasure during treadmill running. Sport Exerc Perform Psychol. 2018;7: 80–92. doi:10.1037/spy0000115

2. Bonnette R, Smith MCI, Spaniol F, Melrose D, Ocker L. The effect of music listening on running performance and rating of percieved exertion of college students. J Strength Cond Res. 2010;24: 1. doi:10.1097/01.JSC.0000367073.45565.4b

3. Karageorghis CI, Priest D-L. Music in the exercise domain: a review and synthesis (Part I). Int Rev Sport Exerc Psychol. 2012;5: 44–66. doi:10.1080/1750984X.2011.631026

4. Terry PC, Karageorghis CI, Saha AM, D’Auria S. Effects of synchronous music on treadmill running among elite triathletes. J Sci Med Sport. 2012;15: 52–57. doi:10.1016/j.jsams.2011.06.003

5. Bacon CJ, Myers TR, Karageorghis CI. Effect of music-movement synchrony on exercise oxygen consumption. J Sports Med Phys Fitness. 2012;52: 359–365.

6. Karageorghis CI, Mouzourides DA, Priest D-L, Sasso TA, Morrish DJ, Walley CJ. Psychophysical and ergogenic effects of synchronous music during treadmill walking. J Sport Exerc Psychol. 2009;31: 18–36. doi:10.1123/jsep.31.1.18

7. Simpson SD, Karageorghis CI. The effects of synchronous music on 400-m sprint performance. J Sports Sci. 2006;24: 1095–1102. doi:10.1080/02640410500432789

8. Bood RJ, Nijssen M, van der Kamp J, Roerdink M. The power of auditory-motor synchronization in sports: enhancing running performance by coupling cadence with the right beats. PloS One. 2013;8: e70758. doi:10.1371/journal.pone.0070758

9. Van Dyck E, Moens B, Buhmann J, Demey M, Coorevits E, Dalla Bella S, et al. Spontaneous entrainment of running cadence to music tempo. Sports Med - Open. 2015;1: 15. doi:10.1186/s40798-015-0025-9

10. Buhmann J, Moens B, Lorenzoni V, Leman M. Shifting the musical beat to influence running cadence. Proc 25th Anniv Conf Eur Soc Cogn Sci Musice. 2017; 27–31.

11. Van Dyck E, Buhmann J, Lorenzoni V. Instructed versus spontaneous entrainment of running cadence to music tempo. Ann N Y Acad Sci. 2021;1489: 91–102. doi:10.1111/nyas.14528

12. Zatorre RJ, Halpern AR, Perry DW, Meyer E, Evans AC. Hearing in the mind’s ear: a pet investigation of musical imagery and perception. J Cogn Neurosci. 1996;8: 29–46. doi:10.1162/jocn.1996.8.1.29

13. Borg GA. Psychophysical bases of perceived exertion. Med Sci Sports Exerc. 1982;14: 377–381.

14. Zemková E, Hamar D. Physiological mechanisms of post-exercise balance impairment. Sports Med Auckl NZ. 2014;44: 437–448. doi:10.1007/s40279-013-0129-7

15. Schütte KH, Maas EA, Exadaktylos V, Berckmans D, Venter RE, Vanwanseele B. Wireless tri-axial trunk accelerometry detects deviations in dynamic center of mass motion due to running-induced fatigue. Triche EW, editor. PLOS ONE. 2015;10: e0141957. doi:10.1371/journal.pone.0141957

16. Faul F, Erdfelder E, Lang A-G, Buchner A. G*Power 3: a flexible statistical power analysis program for the social, behavioral, and biomedical sciences. Behav Res Methods. 2007;39: 175–191. doi:10.3758/bf03193146

17. Centala J, Pogorel C, Pummill SW, Malek MH. Listening to fast-tempo music delays the onset of neuromuscular fatigue. J Strength Cond Res. 2020;34: 617. doi:10.1519/JSC.0000000000003417

18. Fox SM, Naughton JP. Physical activity and the prevention of coronary heart disease. Prev Med. 1972;1: 92–120. doi:10.1016/0091-7435(72)90079-5

19. Jim Valvano. Don’t Give Up. 1993. Available: https://youtu.be/HuoVM9nm42E?feature=shared

20. Edworthy J, Waring H. The effects of music tempo and loudness level on treadmill exercise. Ergonomics. 2006;49: 1597–1610. doi:10.1080/00140130600899104

21. Buhmann J, Moens B, Van Dyck E, Dotov D, Leman M. Optimizing beat synchronized running to music. PloS One. 2018;13: e0208702. doi:10.1371/journal.pone.0208702

22. Brownley KA, McMurray RG, Hackney AC. Effects of music on physiological and affective responses to graded treadmill exercise in trained and untrained runners. Int J Psychophysiol Off J Int Organ Psychophysiol. 1995;19: 193–201. doi:10.1016/0167-8760(95)00007-f

23. Kawabata M, Chua KL. A multiple mediation analysis of the association between asynchronous use of music and running performance. J Sports Sci. 2021;39: 131–137. doi:10.1080/02640414.2020.1809153

24. Clark JC, Baghurst T, Redus BS. Self-selected motivational music on the performance and perceived exertion of runners. J Strength Cond Res. 2021;35: 1656–1661. doi:10.1519/JSC.0000000000002984

25. Lindsay TR, Yaggie JA, McGregor SJ. Contributions of lower extremity kinematics to trunk accelerations during moderate treadmill running. J NeuroEngineering Rehabil. 2014;11. doi:10.1186/1743-0003-11-162

26. Schücker L, Schmeing L, Hagemann N. “Look around while running!” Attentional focus effects in inexperienced runners. Psychol Sport Exerc. 2016;27: 205–212. doi:10.1016/j.psychsport.2016.08.013

27. Limmeroth J, Schücker L, Hagemann N. Don’t stop focusing when it gets harder! The positive effects of focused attention on affective experience at high intensities. J Sports Sci. 2022;40: 2018–2027. doi:10.1080/02640414.2022.2127511

28. Wulf G, McNevin N, Shea CH. The automaticity of complex motor skill learning as a function of attentional focus. Q J Exp Psychol A. 2001;54: 1143–1154. doi:10.1080/713756012

29. Richer N, Saunders D, Polskaia N, Lajoie Y. The effects of attentional focus and cognitive tasks on postural sway may be the result of automaticity. Gait Posture. 2017;54: 45–49. doi:10.1016/j.gaitpost.2017.02.022

